# Independent mechanisms of temporal and linguistic cue correspondence benefiting audiovisual speech processing

**DOI:** 10.1101/2020.07.31.229203

**Authors:** Sara Fiscella, Madeline S Cappelloni, Ross K Maddox

**Affiliations:** Brain and Cognitive Sciences, University of Rochester, Rochester, NY 14627, USA; Del Monte Institute for Neuroscience, University of Rochester, Rochester, NY 14627, USA; Biomedical Engineering, University of Rochester, Rochester, NY 14627, USA; Neuroscience, University of Rochester, Rochester, NY 14627, USA; Center for Visual Science, University of Rochester, Rochester, NY 14627, USA

## Abstract

When listening is difficult, seeing the face of the talker aids speech comprehension. Faces carry both temporal (low-level physical correspondence of mouth movement and auditory speech) and linguistic (learned physical correspondences of mouth shape (viseme) and speech sound (phoneme)) cues. Listeners participated in two experiments investigating how these cues may be used to process sentences when maskers are present. In Experiment I, faces were rotated to disrupt linguistic but not temporal cue correspondence. Listeners suffered a deficit in speech comprehension when the faces were rotated, indicating that visemes are processed in a rotation-dependent manner, and that linguistic cues aid comprehension. In Experiment II, listeners were asked to detect pitch modulation in the target speech with upright and inverted faces that either matched the target or masker speech such that performance differences could be explained by binding, an early multisensory integration mechanism distinct from traditional late integration. Performance in this task replicated previous findings that temporal integration induces binding, but there was no behavioral evidence for a role of linguistic cues in binding. Together these experiments point to temporal cues providing a speech processing benefit through binding and linguistic cues providing a benefit through late integration.

## I. INTRODUCTION

While having a conversation may be easy in quiet environments, noisy environments can render listening a challenging task. Though many of us can listen to our conversation partner with minimal conscious effort, the neural processes by which we accomplish this auditory feat are complex, rely on many stimulus cues, and are poorly understood.

When auditory information is insufficient or difficult to process, visual cues help us listen. Seeing the face of a talker greatly improves our ability to comprehend their speech (Arnold and Hill, 2001; Reisberg et al., 1987). Speaking faces carry both temporal and linguistic cues that are congruent with auditory speech. Specifically, the movements of the talker’s mouth and surrounding areas are temporally coherent with the unique amplitude envelope of the auditory stream of interest, and the shapes the mouth makes are linguistically congruent with the speech sounds produced.

Temporal information is an inherently physical cue and constrained by the dynamics of speech production. The time correlation of mouth movements and speech can help listeners pair the relevant auditory and visual streams (Maddox et al., 2015) or even reduce masking effects of competing auditory streams (Grant and Bernstein, 2019). Linguistic information is provided by the link between specific mouth shapes, called visemes, and the phonemes that they generate. Unlike pure temporal coherence, the link between visemes and phonemes is a learned prior that relies on a listener’s experience with language. The underlying mechanisms that may allow multisensory linguistic information to help us listen are not known.

Many researchers have turned to the McGurk effect to demonstrate the effect of visual linguistic cues on auditory perception. The McGurk effect occurs when observers are concurrently presented with an auditory syllable (e.g. “ba”) and a face which either matches the auditory syllable (e.g. “ba”) or a different syllable (e.g. “ga”). Even though subjects are presented with the same auditory syllable in both visual conditions they often report hearing a fused syllable (e.g. “da”) when the face and auditory speech do not match (Mcgurk and Macdonald, 1976). In order to better understand how the brain processes the linguistic cues associated with the face, studies have looked at the effects of inverting the face. These studies have found that when the face is inverted listeners less accurately identify syllables in the visual alone (“lipreading”) condition, and a higher proportion of people report hearing the auditory syllable than the fused syllable in a multisensory (McGurk) condition (Massaro and Cohen, 1996; Ujiie et al., 2018). Despite the findings that inversion of the face can disrupt the processing that underlies the McGurk effect, the monosyllabic stimuli involved do not well model the demands of listening to speech in noise. It is still unclear whether these linguistic cues carry the same perceptual weight in continuous speech.

Though temporal and linguistic cues both can contribute to speech comprehension, they may contribute in very different ways or at different stages of the multisensory perception process. Multisensory integration has been traditionally thought of as a Bayesian combination of unisensory information just prior to perceptual decision making (“late integration”) (Körding et al., 2007), but recent work has pointed towards an earlier stage of multisensory integration known as “binding” or “early integration” (Atilgan et al., 2018; Bizley et al., 2016; Lee et al., 2019). Binding occurs when an auditory and visual stream are combined by the brain into a single perceptual object. Binding affects the encoding of an audiovisual object, whereas many of the effects of multisensory integration can be explained by a later decision bias (Bizley et al., 2016). When the brain forms a perceptual object, it can then allocate object-based attention (Shinn-Cunningham, 2008). By attending an object, all features are automatically enhanced, even orthogonal features that are not comodulated (Lee et al., 2019). This can both occur within a modality (i.e., multiple visual features combine to form a single visual object (Blaser et al., 2000)) or across modalities (i.e., a visual feature is bound with an auditory feature to form a multisensory object (Maddox et al., 2015)).

Despite potentially having independent neural underpinnings, studies often fail to distinguish between binding and late integration. Binding can be tested in a task that requires the listener to attend two streams to complete a dual task. If the stimuli in the two streams are bound, they can attend to the combined object and improve their performance instead of having to divide their attention to complete the task (Bizley et al., 2016). For binding to occur, there must be some compelling relationship between the object’s features. Possible relationships between features, which may or may not contribute to binding, may roughly be divided into several categories: low-level physical correspondence, semantic congruence, and learned physical correspondence. Low-level physical correspondence involves fundamental relationships of stimuli such as temporal coherence, which is known to induce binding (Maddox et al., 2015), and spatial congruence. Semantic congruence encompasses higher-level learned relationships between stimuli that are commonly paired in the natural world, such as the image and sound of a dog barking, and are unlikely to induce binding due to the significant high-level processing required to make these associations. Learned physical relationships are similar to semantic congruence in the sense that they must be learned through observation of the natural world and similar to low-level physical correspondence in that the physical relationship of the stimuli is inherent to their production, such as visemes and phonemes in which mouth shapes are innately connected to the sounds they produce but are only known to be related by someone with experience of talkers. Here we investigate whether the learned physical relationships of natural speech induce binding.

In Experiment I, we sought to determine if rotating the face of a talker disrupts linguistic cues in a behaviorally relevant task and therefore provide some insight into how these cues contribute to listening in noisy environments. We engaged listeners in a speech in noise task with rotated videos of the target talker to determine how their performance was affected by disrupted cues. We found that rotating the face hindered their speech comprehension, suggesting that the information carried by the face was indeed disrupted by the rotation.

We tested whether linguistic cues could induce binding in Experiment II. Given that the face inversion disrupted linguistic cues, we looked for differences in a multisensory selective attention task that might suggest differences in binding. Here we asked listeners to detect auditory pitch modulations and visual events in target stimuli while ignoring maskers. Binding is a likely mechanism to explain any improvement in performance when the listener saw the target’s face relative to the masker’s face. We found that although there was a clear advantage to seeing the target’s face in all conditions, there was no effect of face rotation, suggesting that the disrupted linguistic cues did not impair binding.

We ultimately find that while linguistic cues are important for listening to speech in noisy environments, this is likely due to late integration rather than binding.

## II. EXPERIMENT I

### A. Methods

Participants performed a speech in noise comprehension task.

#### a. Participants

Fifteen participants (12 female, 3 male) age 18−26 (mean 22.6) had normal hearing (20 dB HL or better at octave frequencies from 500 Hz to 8000 Hz), self-reported normal or corrected-to-normal vision, and spoke English as their primary language. Participants gave written consent and were paid for their participation. All protocols were approved by the University of Rochester Research Subjects Review Board.

#### b. Stimuli

The visual and auditory stimuli were selected from the STeVi corpus. The corpus includes video recordings of four native English speakers (two male and two female) saying 200 high probability sentences that contained three to five keywords each (e.g. “The scarf was made of shiny silk.”) (STeVi Speech Test Video Corpus, n.d.). We rotated the videos by 0°, ±90°, or 180° in each trial. In some trials, still frames from the beginning of the unrotated videos were used instead of the dynamic faces. To ensure that all videos were the same size and aspect ratio, we drew a circle for each trial around the face of the speaker and its dark blue background, leaving the background outside of the circle black. In addition, the mouths of the talkers were centered on the screen (Figure 1). We used high probability sentences due to previous research showing a benefit of semantic context in understanding speech in noisy environments (Van Engen et al., 2014). Due to an error in the experiment code, the videos were played at 29.04 frames per second (fps) instead of their native original 29.97 fps, which led to a small offset that increased throughout the trial with the biggest offset would be at the end of the longest sentence—a delay of 105 ms in the worst case. On average the delay was 37 ms, about one frame of video, and well within the audio-visual temporal binding window (Stevenson et al., 2012).

**Figure 1:**
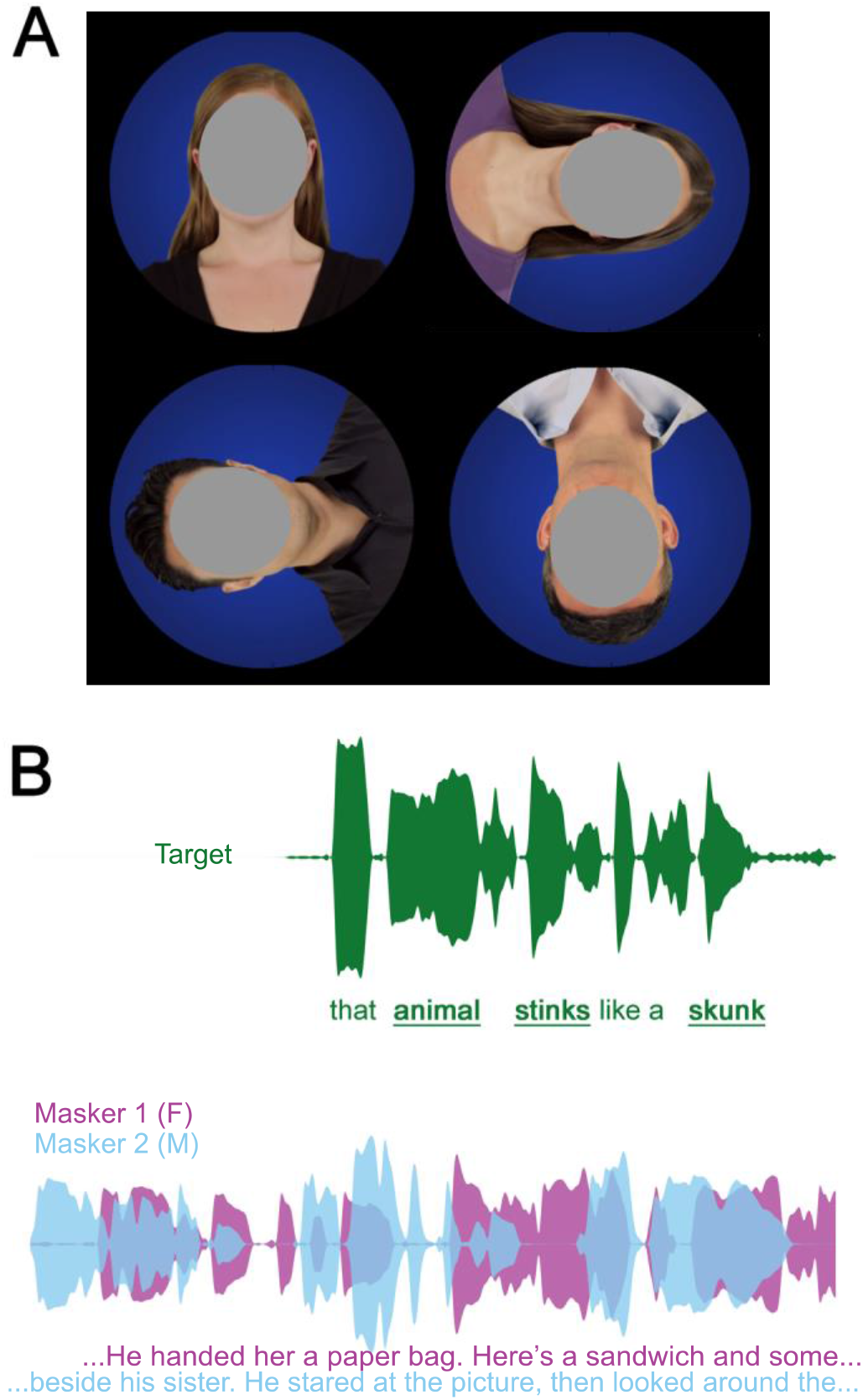
(Color Online). A. Images of the four talkers (faces covered to protect identity of the actors), with 0° (top left), +90° (top right), −90° (lower left), and 180° (lower right) rotation. The same circular mask is applied to all rotations with the mouth centered. B. Envelopes and transcription of auditory stimuli in a given trial. The target sentence began after a 2 second delay (top, green) while two maskers (bottom, female narrator: purple, male narrator: pale blue) played continuously throughout the trial. Subjects had to type each keyword (bold and underlined) in the target sentence while ignoring the maskers in order to receive credit.

There were also two auditory masker streams comprised of natural speech from American English audiobooks, The Alchemyst (Scott, 2008) (male narrator) and A Wrinkle in Time (L’Engle, 2006) (female narrator). Audio was edited to remove silent pauses longer than 0.5 seconds. The masker stimuli were each presented at 60 dB SPL. Then target stimuli were presented with a signal-to-noise ratio (SNR) of 0, −3, or −6 decibels (dB).

#### c. Procedure

Subjects were seated in a dark soundproof booth in front of a 24 inch BenQ monitor, with their nose lined up approximately with the center of the screen and a 50 centimeter viewing distance. Sounds were presented via ER-2 insert earphones (Etymotic Research, Elk Grove Village, Il). Subjects were given a standard keyboard to type in their responses.

Subjects were required to pass a training module before beginning the experiment. Training began with two trials without maskers, followed by three trials of with maskers and an SNR of 0 dB, and lastly three trials with SNR of −3 dB. Participants responded by typing the sentence they heard after each trial with no capitalization or punctuation. For training, where accuracy needed to be judged in real time, responses in a given trial were scored as correct if the sequence of letters of the entered keywords were at least 80% correct. Subjects were given two chances to pass the training, which required correct responses in both of the trials without background noise and two out of the three of the trials with SNR of 0 dB. This served the dual purpose of ensuring that subjects could perform the task and familiarizing them with the talkers’ faces and voices.

Subjects subsequently completed 192 trials. At 25, 50, and 75 percent completion they were given self-timed breaks with a minimum duration of 30 seconds. Each trial consisted of a video of a talker saying a unique high probability sentence with the two background auditory streams playing. No sentences were repeated. The trial began with a 2 second pause on the first frame of the video providing the subject some time to process which voice to listen to for that trial. There were 12 randomly interleaved conditions: three SNRs (0, −3, and −6 dB) and four visual conditions (rotation of 0°, ±90°, or 180°, and a static upright image). After the video played, subjects were instructed to type in what they heard with minimal spelling errors. They were also informed that they would receive partial credit and to give a best guess if they were not certain.

#### f. Scoring

The sentences from the STeVi corpus had three to five predetermined keywords per sentence. The percent of accurately entered keywords was hand scored based on Smayda et al’s (2016) criteria. Responses were considered correct if the spelling errors did not change the meaning of the word or if the words were homophones.

#### g. Statistics

We fit the data to a linear mixed effects model considering both SNR and angle to be categorical variables (we did not expect a linear relationship, nor could we assume monotonicity, particularly with regards to angle). We also considered interactions of SNR and angle. Each subject was fit with an intercept.

### B. Results

In the static condition, in which the face gives neither temporal nor linguistic information, subjects had near chance performance in the speech in noise task at negative signal to noise ratios (Figure 1A). Only at 0 dB SNR did subjects perform reliably above chance for the static face. Across SNRs, subjects were able to significantly improve their performance when the face was moving by 7–47%, depending on the SNR and face rotation (Figure 1B). Across face rotation conditions these gains were largest at 0 dB SNR and decreased as SNR worsened. Within each SNR, improvements were largest for the upright face (0° rotation), and smallest for the inverted face (180° rotation).

Averaging across SNRs to specifically investigate the effect of disrupted linguistic cues, there were significant differences in each subject’s improvement depending on the rotation of the face. The mixed effects model showed that 0 dB and −3 dB conditions were significantly different (p=1.15×10^−10^), but there was not a difference between −3 dB and −6 dB. The 0° and 180° rotations were significantly different(p=7.71×10^−3^), as were the 90° and 180° rotations (p=0.0119). The difference between 0° and 90° rotations approached, but did not reach, significance (p=0.0876). There were no significant interaction terms.

### C. Summary

In Experiment I we demonstrated that rotation of the head disrupted speech comprehension, suggesting orientation specific processing of the face. Given the significant reduction in speech processing between upright (0°) and inverted (180°) faces, we used these rotations to probe the question of whether binding is affected by linguistic cues in Experiment II.

## III. EXPERIMENT II

### A. Methods

We engaged participants in a fundamental frequency modulation discrimination task. Listeners were asked whether a target talker was modulated in pitch while ignoring masker talkers. They simultaneously performed a visual detection catch trial task to ensure visual attention was maintained throughout the experiment.

#### a. Participants

23 participants (17 female, 6 male) ages 19−35 (mean 23.1) met the same criteria as the participants from Experiment I. Six of the participants had also participated in Experiment I and had similar performance to those who were naïve to the stimuli.

#### b. Stimuli

Target and masker high context sentences were selected from the STeVi corpus. The average duration of these sentences was 2.36 seconds with a standard deviation of 0.28 seconds. Each trial included two of these sentences (one target and one masker sentence). These voice pairings were evenly distributed across trials and always consisted of one male and one female talker. The sentence pairings were randomly chosen, with each target sentence presented twice, to have similar durations. All but nine sentence pairs had duration differences under 100 ms and the maximum duration difference was 274 ms.

For some trials the audio from the videos was pitch modulated. Pitch modulations were 10 Hz cosine modulations with peak-to-peak amplitude of two semitones added to the stimulus’ natural pitch trajectory using Praat (Boesma and Weenick, n.d.). Videos were presented upright or were inverted (rotated 180 degrees). The same audiobooks from Experiment I were played at −6 dB SPL to provide additional interfering speech noise and make the task appropriately challenging.

#### c. Procedure

Subjects were given three chances to pass a training module in which they had to detect pitch modulation when no maskers were present. As in Experiment I, this allowed subjects to learn the identities of the talkers. They were given ten practice trials and ten testing trials for which they had to achieve 70% accuracy to pass. The statistics of modulation were the same for both the training and the main experiment. Subjects were then shown two example trials to familiarize them with the complexity of the stimulus. They were instructed to look at the faces on the screen and perform two tasks simultaneously: the main pitch modulation discrimination task, and a catch trial visual detection task. Breaks were offered as in Experiment I.

#### d. Main Task

For each trial there was a target talker and masker talker. At the beginning of the trial, the subject saw an image of the target talker for 1.5 seconds, indicating which to listen to during the trial. After the image was presented, the video and four auditory streams were played: the target talker, the masker talker, and the two interfering audiobook background streams. Subjects reported whether the voice of the target talker contained the modulation by pressing a button.

This task consisted of 16 conditions (2 masker/target video × 2 face rotations × 4 masker/target modulations). The video of a talker either matched the target or the masker auditory stream, the face of the talker was upright or inverted, and both, neither, only the target, or only the masker auditory streams were modulated. The subjects heard each of the 200 high probability sentences twice as a target, for a total of 400 trials. There were 35 trials for each condition in which only the masker or target were modulated (70% of trials; the conditions of interest) and 15 trials for remaining conditions which were included so the subject could not infer the target modulation based on the masker modulation.

#### e. Catch Trials

Of the 120 trails for which either both or neither sentences were modulated, 36 random trials had a small pink translucent dot over the mouth of the talker. Subjects were instructed to press 3 when they saw this dot and to not respond for pitch modulation. These catch trials ensured that subjects were looking at the talker’s face. Subjects were informed each time they failed to detect the dot. The criterion to be included in the analysis was detection of more than 80% of the dots. All subjects achieved this, so all data were included in the analysis.

### B. Results

All subjects were able to perform the pitch discrimination task well above chance (Figure 2A). There was a significant improvement in both the upright (paired t-test, t=6.61, p=1.20×10^−6^) and inverted (paired t-test, t=3.91, p=7.49×10^−4^) conditions when subjects viewed the target face relative to when they viewed the masker face (Figure 2B). However, there was no difference in the benefit due to the target face between the upright and inverted condition, and therefore no benefit of the upright face (Figure 2C). Averaging across face rotation conditions, subjects experienced a significant benefit (paired t-test, t=6.61, p=1.20×10^−6^) of approximately 7% when the face of the video matched the target rather than the masker.

**Figure 2:**
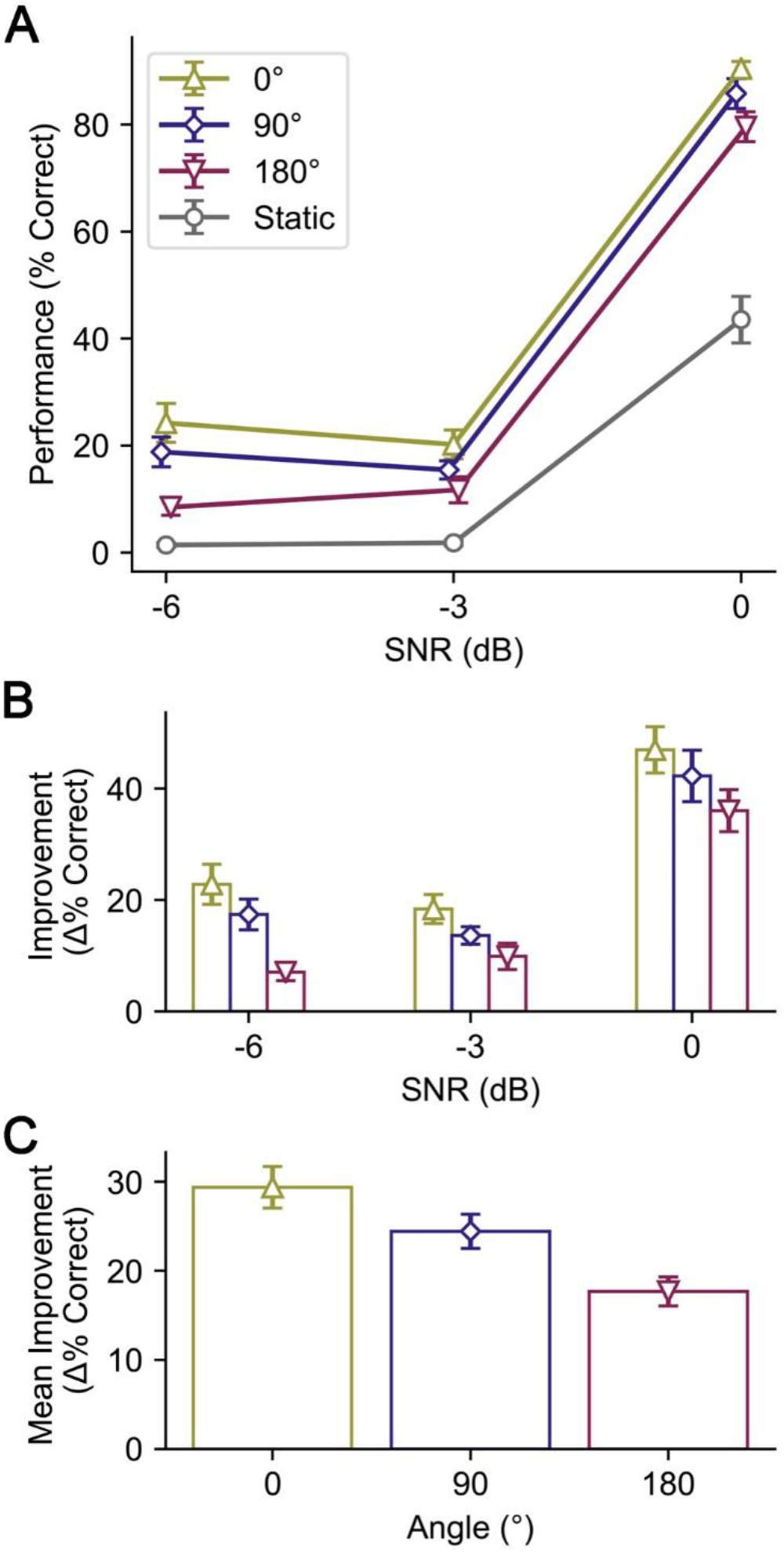
(Color Online). A. Performance in the speech in noise task averaged across subjects. Upward-facing triangle indicates the unrotated or upright face, diamond indicates rotation of the face by 90° to the left or right, downward-facing triangle indicated the inverted face, circle indicates a static image. B. The improvement in performance due to the moving face calculated as the difference in each video condition and the respective static face for a given SNR. C. Performance improvement due to temporal and linguistic cues averaged across SNR conditions. All error bars show ±1 SEM.

**Figure 3:**
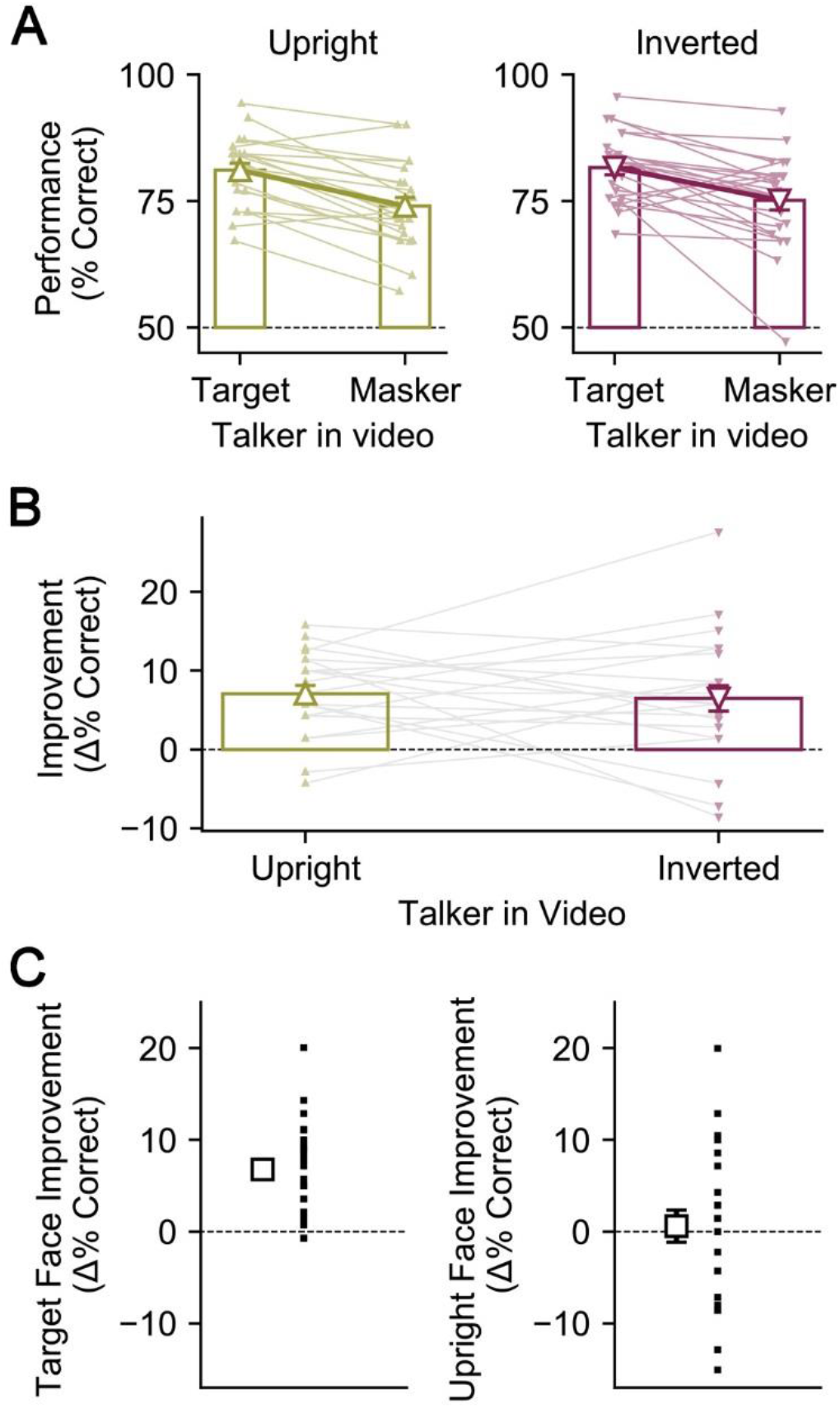
(Color Online). A. Performance in the pitch modulation discrimination task. Small solid markers show individual subjects, larger open markers show the average across subjects. B. The improvement in performance due to the video of the target talker’s face calculated as the difference between the target visual condition and masker visual condition for a given face rotation. C. (Left) Net improvement due to the target talker’s face (difference between target and masker conditions averaged across subjects and upright/inverted conditions). (Right) Net improvement due to the upright face (difference between upright and inverted conditions averaged across subjects and target/masker conditions). All error bars show ±1 SEM.

### C. Summary

In Experiment II the rotation of the face did not significantly influence binding. Nonetheless this paradigm showed a strong replication of previous findings that temporal coherence induces binding (Atilgan et al., 2018; Maddox et al., 2015). The importance of temporal coherence for binding has not previously been established for speech.

## IV. DISCUSSION

Together these experiments suggest that visual linguistic cues and audio-visual binding contribute independently to processing multimodal speech in noise. Experiment I addressed the benefit of visual linguistic cues in speech comprehension and Experiment II investigated audio-visual binding, ultimately showing that visual linguistic cues do not enhance the listener’s object-based attention to the target talker.

In Experiment I we demonstrated that both temporal and linguistic cues are important for speech comprehension in a speech in noise task. There was a significant improvement in performance when the face was moving relative to the static image. This was true of all face rotation conditions. Because temporal cues were preserved across rotation conditions, we consider some portion of the video performance improvement to be due to temporal cues. However, as the rotation of the face increased in magnitude, the benefit of the video decreased, suggesting that some of the benefit in each condition is due to processing of linguistic cue. Though linguistic cues are present even in the rotated faces, the processing of this information seems to be impaired by rotating the face. Interestingly, performance drops with the magnitude of the rotation even though the 0° and 180° rotation are more geometrically similar due to their vertical symmetry than the 0° and 90° rotations. There are two possible explanations for this: the subject can partially compensate for the face’s rotation when processing visemes or that the subject has more prior experience with 90° rotated faces than with 180° rotated faces. While the latter is likely true, we do not believe it is a compelling explanation for our results. A vast majority of conversations are held with upright faces, and situations in which we are speaking to someone at a 90° rotation are minimal (e.g. talking to someone while reclined). Therefore, it seems more likely that subjects are “un-rotating” the face where possible to get some benefit from linguistic cues, and this is easier for them to do with 90° rotation than 180°.

In Experiment II we show that fundamental frequency modulation discrimination is improved when listeners can see the video of the target talker rather than a masker talker regardless of the orientation of the face. Structurally our experiment was very similar to previous work that tested for binding by engaging listeners in simultaneous auditory discrimination and visual detection tasks (Maddox et al., 2015). Importantly, the tasks rely on the tracking of an orthogonal perceptual feature (pitch), one that is independently changing to the feature that is coherently modulated. If the listener binds the auditory and visual streams based on their temporal coherence, their brain will form a perceptual object. By allocating object-based attention, all features of the object, including the orthogonal feature will be enhanced, leading to better performance. The performance improvement is not explained by late-stage integration since the visual stream provides no information about the orthogonal auditory features.

We improved upon Maddox et al’s original task (2015) by using more natural stimuli. In this case the listeners had to simultaneously determine whether there was a pitch modulation in the target talker or a pink disk on the mouth in the video while ignoring the masker talker. We used real speech as the stimuli and pitch as the orthogonal feature, which gave ecological relevance to the task. Processing of pitch modulations are important in natural environments due to prosodic information that is in part carried by the pitch of a talker. This prosodic information not only provides emotional context but also is important for parsing full sentences (Stirling, 1996; Warren et al., 1995). Binding of audiovisual speech could improve our perception of not only what the talker is saying, but how they are saying it. Binding can explain an improvement in performance when the video matches the target. The listener can allocate object-based attention to the target talker and improve their discrimination of pitch modulation because detection of visual events will not divide their attention.

There is a consistent improvement in processing orthogonal stimulus features when the listener can see the target video, which can be explained by binding. However, the benefit is not modulated by rotating the face, suggesting that temporal coherence is the cue that underpins binding in this experiment. Using real speech, our results confirm the finding that temporal coherence drives audiovisual binding, which had been previously established for stimuli with speech-like dynamics (Maddox et al., 2015).

We did not find an effect of face rotation on performance in the pitch discrimination task. There are a few possible explanations for this. Temporal coherence may be sufficient to induce binding, and a possible contribution of linguistic cues would be overshadowed by the influence of temporally coherent cues. Alternatively, a contribution of linguistic cues to strengthen binding, if such a thing is possible, may have been too small to be measured behaviorally. If linguistic cues truly do not influence binding, the hierarchical processing of language therefore suggests an explanation for binding occurring independent of face rotation. While low level spectral features are well represented in A1, articulatory features are not represented until the superior temporal gyrus (STG) (Ding et al., 2016; Mesgarani et al., 2014). A study involving ferrets performing a multisensory task found neural evidence of binding in primary auditory cortex (A1) (Atilgan et al., 2018), whereas traditional Bayesian or late-stage integration is thought to occur at higher processing areas in the intraparietal sulcus (Rohe and Noppeney, 2015, 2016). Such findings support the notion that binding and late-stage integration are fundamentally different processes that rely on different types of sensory information. The extent of the visual and auditory information available at such early processing areas to create binding is uncertain, particularly given the unknown origins of the visual connections responsible for visual-dependent auditory activity in A1. In order for linguistic cues to contribute to binding the brain would need to combine feedback from STG carrying auditory articulatory information with viseme information from visual areas.

Together these experiments show that face-rotation and therefore disruption of linguistic cues hinders audiovisual speech comprehension, but not detection of orthogonal pitch features. Even if linguistic cues do play a role in binding, their behavioral benefit seems to be superseded by temporal coherence. Therefore, the benefit of visual linguistic cues to speech understanding is likely due to late integration in which visemes can bias the listener towards the correct phoneme perception at higher processing stages. Binding, then, may be specific to very low-level physical correspondences, a hypothesis on which future experiments will shed more light.

## V. CONCLUSION

We demonstrated the importance of both temporal and linguistic visual cues for audiovisual speech in noise comprehension in an ecologically relevant task. We also showed that audiovisual temporal coherence, but not linguistic congruence, improved performance in a frequency modulation discrimination task, consistent with the existence of audiovisual binding. It thus appears that multisensory linguistic cues, an example of learned physical correspondence, are integrated at the perceptual decision-making stage rather than early integration. Practically, our results suggest that visemes can benefit listeners in noisy environments by biasing the listener towards perceiving the correct sentence, but they do not aid listeners in detecting other aspects of the talker’s speech.

## ACKNOWLEDGMENTS

Research reported in this publication was supported by the National Institute on Deafness and Other Communication Disorders of the National Institutes of Health under award number R00DC014288.

## REFERENCES

Arnold, P., and Hill, F. (2001). “Bisensory augmentation: A speechreading advantage when speech is clearly audible and intact,” Br. J. Psychol., 92, 339–355. doi:10.1348/000712601162220

Atilgan, H., Town, S. M., Wood, K. C., Jones, G. P., Maddox, R. K., Lee, A. K. C., and Bizley, J. K. (2018). “Integration of Visual Information in Auditory Cortex Promotes Auditory Scene Analysis through Multisensory Binding,” Neuron, 97, 640–655.e4. doi:10.1016/j.neuron.2017.12.034

Bizley, J. K., Maddox, R. K., and Lee, A. K. C. (2016). “Defining Auditory-Visual Objects: Behavioral Tests and Physiological Mechanisms,” Trends Neurosci., 39, 74–85. doi:10.1016/j.tins.2015.12.007

Blaser, E., Pylyshyn, Z. W., and Holcombe, A. O. (2000). “Tracking an object through feature space,” Nature, 408, 196–.

Boesma, P., and Weenick, D. (n.d.). Praat: doing phonetics by computer,.

Ding, N., Melloni, L., Zhang, H., Tian, X., and Poeppel, D. (2016). “Cortical tracking of hierarchical linguistic structures in connected speech,” Nat. Neurosci., 19, 158–164. doi:10.1038/nn.4186

Grant, K. W., and Bernstein, J. G. W. (2019). “Toward a Model of Auditory-Visual Speech Intelligibility,” In A. K. C. Lee, M. T. Wallace, A. B. Coffin, A. N. Popper, and R. R. Fay (Eds.), Multisensory Process. Audit. Perspect., Springer Handbook of Auditory Research, Springer International Publishing, Cham, pp. 33–57. doi:10.1007/978-3-030-10461-0_3

Körding, K. P., Beierholm, U., Ma, W. J., Quartz, S., Tenenbaum, J. B., and Shams, L. (2007). “Causal Inference in Multisensory Perception,” PLOS ONE, 2, e943. doi:10.1371/journal.pone.0000943

Lee, A. K. C., Maddox, R. K., and Bizley, J. K. (2019). “An Object-Based Interpretation of Audiovisual Processing,” In A. K. C. Lee, M. T. Wallace, A. B. Coffin, A. N. Popper, and R. R. Fay (Eds.), Multisensory Process. Audit. Perspect., Springer Handbook of Auditory Research, Springer International Publishing, Cham, pp. 59–83. doi:10.1007/978-3-030-10461-0_4

L’Engle, M. (2006). A Wrinkle in Time, Findaway World Llc, 238 pages.

Maddox, R. K., Atilgan, H., Bizley, J. K., and Lee, A. K. (2015). “Auditory selective attention is enhanced by a task-irrelevant temporally coherent visual stimulus in human listeners,” eLife, , doi: 10.7554/eLife.04995. doi:10.7554/eLife.04995

Massaro, D. W., and Cohen, M. M. (1996). “Perceiving speech from inverted faces,” Percept. Psychophys., 58, 1047–1065. doi:10.3758/BF03206832

Mcgurk, H., and Macdonald, J. (1976). “Hearing lips and seeing voices,” Nature, 264, 746–748. doi:10.1038/264746a0

Mesgarani, N., Cheung, C., Johnson, K., and Chang, E. F. (2014). “Phonetic Feature Encoding in Human Superior Temporal Gyrus,” Science, 343, 1006–1010. doi:10.1126/science.1245994

Reisberg, D., McLean, J., and Goldfield, A. (1987). “Easy to hear but hard to understand: A lip-reading advantage with intact auditory stimuli,” Hear. Eye Psychol. Lip-Read., Lawrence Erlbaum Associates, Inc, Hillsdale, NJ, US, pp. 97–113.

Rohe, T., and Noppeney, U. (2015). “Cortical Hierarchies Perform Bayesian Causal Inference in Multisensory Perception,” (C. Kayser, Ed.) PLOS Biol., 13, e1002073. doi:10.1371/journal.pbio.1002073

Rohe, T., and Noppeney, U. (2016). “Distinct Computational Principles Govern Multisensory Integration in Primary Sensory and Association Cortices,” Curr. Biol., 26, 509–514. doi:10.1016/j.cub.2015.12.056

Scott, M. (2008). The Alchemyst: The Secrets of the Immortal Nicholas Flamel, Delacorte Press, 402 pages.

Shinn-Cunningham, B. G. (2008). “Object-based auditory and visual attention,” Trends Cogn. Sci., 12, 182–186. doi:10.1016/j.tics.2008.02.003

Smayda, K. E., Engen, K. J. V., Maddox, W. T., and Chandrasekaran, B. (2016). “Audio-Visual and Meaningful Semantic Context Enhancements in Older and Younger Adults,” PLOS ONE, 11, e0152773. doi:10.1371/journal.pone.0152773

Stevenson, R. A., Zemtsov, R. K., and Wallace, M. T. (2012). “Individual Differences in the Multisensory Temporal Binding Window Predict Susceptibility to Audiovisual Illusions,” J. Exp. Psychol. Hum. Percept. Perform., 38, 1517–1529. doi:10.1037/a0027339

STeVi Speech Test Video Corpus (n.d.). Sensimetrics. Retrieved from https://www.sens.com/products/stevi-speech-test-video-corpus/

Stirling, L. (1996). “Does Prosody Support or Direct Sentence Processing?,” Lang. Cogn. Process., 11, 193–212. doi:10.1080/016909696387268

Ujiie, Y., Asai, T., and Wakabayashi, A. (2018). “Individual differences and the effect of face configuration information in the McGurk effect,” Exp. Brain Res., 236, 973–984. doi:10.1007/s00221-018-5188-4

Van Engen, K. J., Phelps, J. E. B., Smiljanic, R., and Chandrasekaran, B. (2014). “Enhancing speech intelligibility: interactions among context, modality, speech style, and masker,” J. Speech Lang. Hear. Res. JSLHR, 57, 1908–1918. doi:10.1044/JSLHR-H-13-0076

Warren, P., Grabe, E., and Nolan, F. (1995). “Prosody, phonology and parsing in closure ambiguities,” Lang. Cogn. Process., 10, 457–486. doi:10.1080/01690969508407112

